# Preclinical study of the human recombinant Erythropoietin (GBPD002) compared with Eprex^®^

**DOI:** 10.1101/2024.06.05.597513

**Authors:** Kakon Nag, Mohammad Mohiuddin, Maksudur Rahman Khan, Samir Kumar, Enamul Haq Sarker, Rony Roy, Bipul Kumar Biswas, Uttam Barman, Sheik Rejaul Haq, Sitesh Chandra Bachar, Naznin Sultana

## Abstract

Erythropoietin (EPO) is a glycoprotein that an essential growth factor for erythroid progenitors at the bone marrow, which appears in blood as a response to hypoxia. It is produced mainly by the kidneys; and its biosynthesis and release are stimulated by the reduction of tissue oxygenation and/or the reduction of the mass of erythrocytes. Here, we report the results of the preclinical evaluation of the safety of ‘GBPD002’ a recombinant human erythropoietin (rhEPO) developed by the Globe Biotech Limited, Bangladesh, through a comparative study of acute toxicity with Eprex^®^, a commercial homologue from Janssen, UK. The product was administered subcutaneously into Wistar rats, at 500 lU/kg of therapeutic dose (TD) and 3 times of TD for the single dose toxicity study on 14 consecutive days and 125 UL/kg, 250 UL/kg, 500 UL/kg, 750 UL/kg, 1000 UL/kg, 1250 UL/kg and 1500 UL/kg of GBPD002 and Eprex^®^ on 7 consecutive days respectively for the repeated dose toxicity study. Hematological and biochemical parameters were measured for all test subjects before first dose injection and the day after last dose injection of the both studies. Necropsy and histopathology of representative subjects from each group were also observed to find any pathological significance like degeneration or cellular necrosis in internal organs such as kidney, liver, lung and spleen of any rat under experiment. Both GBPD002 and Eprex^®^ comparative toxicology studies revealed similar pharmacologically driven mechanisms of toxicity, which is statistically insignificant (*p* >0.05). Though hematology parameter values stayed within the normal range during the assay period but the high count of hemoglobin and high hematocrit (*P*<0.05), together with the decrease in white blood cell, confirm the therapeutic effect of Erythropoietin in both studies. Moreover, in both studies, the initial and final values of aspartate aminotransferase, alanine aminotransferase and blood urea nitrogen were also found similar (*p* >0.05) for both GBPD002 and Eprex^®^ in the study. The study clearly established that the toxicological profile of “GBPD002” and Eprex^®^, administered subcutaneously, were similar and related to the known pharmacology of erythropoietin alfa; hereby, demonstrating the proof of ‘totality’ and ‘no residual uncertainty’ between “GBPD002” and Eprex^®^. Therefore, “GBPD002” and Eprex^®^ shall be administered interchangeably in relevant indications.

## 1. Introduction

The hormone erythropoietin (EPO), a glycoprotein, is essential for the production of red blood cells (RBCs). EPO is mostly produced in the kidney’s peritubular cells and released into the bloodstream in adults [1]. The EPO receptor on bone marrow erythroid progenitors is bound by circulating EPO, initiating a number of signaling pathways that promote the development of mature RBCs [2]. It was originally isolated from the urine of aplastic anemia patients and characterized as a 34 000 Da protein [3]. The expression of this endogenous glycoprotein is controlled by the transcription factor, hypoxia-inducible factor [4]. 40% of the molecule is made up of carbohydrates, which are primarily sialic acid and are dispersed at three N-linked and one O-linked glycosylation site. These carbs are necessary for EPO to function biologically [5]. The molecule’s terminal sialic acid residues prevent it from being quickly cleared by the liver, preserving its level of activity in the bone marrow. Conversely, the biological activity of EPO in vivo is lost when sialic acid is eliminated enzymatically by neuraminidases [6]. The physiological site of EPO synthesis is the kidneys, and a specific reduction in the oxygen supply to the kidneys or broad anemia both cause an increase in EPO production. Via certain receptors on the cell surface, EPO interacts with its target cells. These receptors are expressed mostly at erythroid progenitor cells, but they are also present in neuronal, endothelial, multipotent hematopoietic, and embryonic stem cells [7].

As a medicinal treatment, erythropoietin is made using recombinant DNA technology in mammalian cell cultures that have been transfected with the human EPO gene [8]. Recombinant DNA technology has enabled the production of human EPO (rhEPO), allowing its extensive therapeutic application in clinical settings [9]. A Note on Guidance appeared and provided recommendations mainly for non-clinical tests to be performed following changes in manufacture of a biologic, but also for a product claimed to be similar to another one already authorized of the data protection period [10, 11]. A series of in vitro studies on receptor binding and in vivo investigations were required under the guidelines “if there are specific uncertainties or concerns regarding safety” [12]. Clinical evidence suggests that it can be used to treat anemia associated to entities like chronic renal insufficiencies, rheumatoid arthritis, premature birth, chemotherapy, transfusions and oncohematological diseases, among others [13]. When EPO is administered, no allergies have been reported; nevertheless, a small number of patients have had arthralgias and local cutaneous responses [14]. Preclinical trials involving EPO are important for establishing its safety, efficacy, and understanding its potential therapeutic applications across various medical conditions [15].

This manuscript purpose to provide a comprehensive overview of the preclinical studies conducted on human recombinant erythropoietin, with a focus on its safety, efficacy, and potential as a drug product. As a prelude to clinical trials and subsequent regulatory approval, preclinical investigations play a pivotal role in elucidating the pharmacological properties, toxicological profiles, and mechanisms of action of novel therapeutic agents [16, 17]. The exploration of rhEPO begins with an examination of the protein’s structure and the underlying genetic modifications that render it a recombinant biopharmaceutical [18]. By delving into the molecular intricacies of rhEPO, we aim to establish a foundational understanding of its design and functionality. Additionally, this section will highlight the significance of rhEPO in the context of endogenous erythropoietin, emphasizing the potential advantages and limitations of using a recombinant form.

The first-ever alpha epoetin product, Eprex^®^, is a prescription medication with a track record of safety and effectiveness [19]. Eprex^®^was manufactured by Johnson & Johnson and it was the first EPO formulation to receive regulatory approval in Europe in 1988. In the early 1990s, physicians outside of the United States adopted the subcutaneous route of administration of EPO for hemodialysis patients due to the socio-economic benefit for the patients [20]. GBPD002 is a biosimilar of Eprex^®^, which is developed by Globe Biotech Limited and synthesized in genetically engineered Chinese hamster ovary (CHO) cells. Upstream and downstream process development and validation were done for large-scale production [21]. Step-by-step analytical results confirmed the biosimilarity of GBPD002 with Eprex^®^ regarding molecular characterization [22].

Moving forward, the manuscript will delve into the preclinical safety assessments, encompassing acute and chronic toxicity studies in relevant animal models. Comprehensive evaluations of potential adverse effects, dose-dependent responses, and organ-specific toxicity will be crucial in establishing a safety profile for rhEPO, guiding subsequent clinical trial designs and mitigating risks associated with human administration [23]. A study designed for the preclinical assessment of the safety of GBPD002 is presented here that includes the evaluation of single and repeated dose toxicity. With this goal, the product was compared to a commercial homologue (Eprex^®^) for the evaluation of single and repeat-dose toxicity were performed in Wister rats to analyze the toxicity of GBPD002.

## 2. Materials and Methods

The studies were performed in compliance with the ethical guidelines for the use of animals in experimentation, Good Laboratory Practices and approved Standard Operating Procedures for the implementation of toxicological studies.

### 2.1 Formulation

The sample was formulated using formulation buffer and AKTA flux s (GE Healthcare, USA). After sterile filtration through 0.22-micron PES Sartopore 2 filter (Sartorius Stedim, Germany), pre-formulated bulk sample was transferred to quality control (QC) for testing as per specification. Then, sterile and pre-formulated bulk drug substance in 2 D single use bag (Sartorius Stedim, France) was transferred to fill-finish facility for filling & packaging maintaining 4 – 8°C temperature. The sterile 1 ml empty long syringes (Schott, Switzerland) were filled using automatic combo filling and closing machine (Tofflon, China) at 1 ml volume for dose preparation. The pre-filled syringes were packaged (blistering) using blister machine HM-AV plus (Hoonga, The Republic of Korea). After blistering completion, the drug product (GBPD002) was stored at 4 – 8°C. Finally, the pre-filled syringes were tested by quality control as per specification and transferred to pre-clinical (animal) study center maintaining 4 – 8°C temperature for toxicology study.

### 2.2 Animal selection

Wistar rats (*Rattus norvegicus*) were used for the single and repeated dose safety and toxicity study, respectively. A total number of 75 Wistar male and female rats of 10 – 25 weeks old were selected and isolated 5 days before the dosing. After attentive monitoring and conditioning, 36 rats (18 males and 18 females) were subjected for single dose toxicity analysis and alternately, 24 rats (12 males and 12 females) were subjected for repeat dose toxicity analysis. The temperature of the experimental animal room was 26 °C (±2 °C) and the relative humidity was 60±5%. The room was HVAC controlled ISO class 7 room with 70% fresh air intake and full exhaust. The rats were individually housed in polypropylene cage with proper water and feed, and kept under 12 hours of day-night cycle. Target weight of male animals was 185±20 g and female animals were 175±20 g. In the entire experiment, 20% additional animals (males and females) were use as substitutes for the excluded animals. Healthy young adult animals were often used in laboratory experiments. Females were nulliparous and non-pregnant. At the outset of the study, each animal was between 10 and 15 weeks old. The weight difference of animals used was also minimal and not exceeding ±20% of the mean weight of each sex. These animals were used to return any individuals kept out during the study periods. The study plan and procedures were approved by the internal ethical review board of Globe Biotech Limited, which is complied with local ethical regulation. No treatment randomization and blinding methods were used in the study and sample sizes were determined by the resource equation method.

### 2.3 Experimental Design

#### 2.3.1 Single dose toxicity assay

A total number of 36 Wister rats including both male and female were used for the study comprising 4 treatment groups, 1 placebo group and 1 control group where each group consists of 3 male and 3 female rats. Among the 4 treatment groups, 2 groups were assigned for normal (500 IU/kg) and toxic dose (1500 IU/kg) respectively, to support the toxicological similarity of GBPD002 to Eprex^®^. Treatment-1 and Treatment-2 were injected with normal and toxic dose of GBPD002, respectively. Similarly, Treatment-3 and Treatment-4 were injected with normal and toxic dose of Eprex^®^, respectively. The placebo group was assigned to inject 100 µL of formulation buffer for GBPD002 and Eprex^®^. The control group was assigned without any injection as negative control of the study. Pre-dose whole blood (approximately 200 µL) from each rat was collected for complete blood count (CBC) in 2% EDTA at 2 days before dosing. Similarly, whole blood was also collected after last dosing at day 14 for CBC analysis. PD endpoints including red blood cell count (RBC), white blood cell count (WBC), hemoglobin (HGB), hematocrit (HCT), mean corpuscular volume (MCV), mean corpuscular hemoglobin (MCH) and mean corpuscular hemoglobin concentration (MCHC) were measured for all test subjects before injection and the last day of 14 days’ study using auto hematology analyzer BK-6190-Vet (Biobase, Germany). Similarly, Alanine aminotransferase (ALT). Aspartate aminotransferase (AST), and Blood urea nitrogen (BUN) assay were also performed from blood serum using semi-automatic chemistry analyzer (Biobase, Germany) for checking the toxic effect of GBPD002 and Eprex^®^ on liver and kidney. Body temperature and weight also measured during the whole study period. Necropsy of representative subject animals from each treatment, placebo and control group were done to check for any abnormalities in kidney, lung, liver, spleen and heart. External surfaces, all orifices, cranial cavities, external surfaces of brain and spinal cord, thoracic abdominal and pelvic cavities, cervical tissues and organs were also observed during necropsy test. Finally, histopathology was also done to find any pathological significance like degeneration or cellular necrosis in internal organs such as kidney, liver, lung and spleen of any rat under experiment.

#### 2.3.1 Repeat dose toxicity assay

A total number of 24 rats were separated into 4 different groups consisting 6 rats (3 males and 3 females) in each group. There were 2 different treatment groups such as Treatment group 1 and 2, 1 placebo group and 1 control group. Each rat of treatment group 1 and group 2 was subcutaneously (SC) injected with sterile 125 IU/kg, 250 IU /kg, 500 IU /kg, 750 IU /kg, 1000 IU /kg, 1250 IU /kg and 1500 IU /kg of GBPD002 and Eprex^®^ on 7 consecutive days, respectively. Each rat of the placebo group was injected with 100 µL of vehicle on 7 consecutive days and the control group was assigned without any injection as negative control of the study. Pre-dose whole blood (approximately 200 µL) from each rat was collected for complete CBC in 2% EDTA at 2 days before dosing. Similarly, whole blood was also collected after last dosing at day 7 for CBC analysis using auto hematology analyzer BK-6190-Vet (Biobase, Germany). PD endpoints including RBC, WBC, HGB, HCT, MCV, MCH, MCHC and Platelet (PLT) were measured for all test subjects before first dose injection and the day after last dose. Similarly, ALT, AST, and BUN assay were also performed similarly from blood serum using semi-automatic chemistry analyzer (Biobase, Germany) for checking the toxic effect of GBPD002 and Eprex^®^ on liver and kidney. Body temperature and weight also measured on the whole study period. Necropsy of representative subject animals from each treatment, placebo and control group were done to check for any abnormalities in kidney, lung, liver, spleen and heart. External surfaces, all orifices, cranial cavities, external surfaces of brain and spinal cord, thoracic abdominal and pelvic cavities, cervical tissues and organs were observed during necropsy test. Finally, histopathology was also done to find any pathological significance like degeneration or cellular necrosis in internal organs such as kidney, liver, lung and spleen of any rat under experiment.

### 2.4 Data Evaluation and Statistical Analysis

The variables that will be used for statistical processing are body weight (BW), body temperature (BT), the hematological and biochemical parameters and the microscopy findings. In all cases, central tendency and dispersion statistics will be calculated (mean, standard deviation, maximum and minimum values). For the treatment of the hematological and biochemical parameters, the variable FD (difference between the final and initial value) will be created. The assumptions of normal distribution and homogeneity of variance will be verified by the Kolmogorov Smirnov and Shapiro-Wilk tests and by the Levene test, respectively, before the analysis of the BW and BT variables for each evaluation time point. Depending on whether the data fit a normal distribution or not, a parametric analysis of variance (ANOVA) or a non-parametric alternative (Kruskall-Wallis test) will be used. Paired comparisons will also be performed on consecutive intervals, using either the paired t-test or the Wilcoxon test, depending on whether the data fit a normal distribution. The results from the histopathological studies will be analyzed by making cross-tabulated classification tables, using the test for associated independence (Fisher’s exact test). The data will be processed with the Microsoft Excel, 2010, running on the Windows operating system. The mean difference between the test and the comparator product was calculated using the linear mixed-effect analysis of variance, along with *p*-value. Statistical significance was defined as a *p*-value of less than 0.05; the *p*-value for PD marker for the sample formulations greater than 0.05 were considered similar and non-significant.

## 3. Results

### 3.1 Body weight

Body weight increased steadily and significantly during the single dose toxicity study, as shown in **figure 1 (A)**. Those differences of body weight in all study groups in different evaluation time point (day 0, 7 and 14), the body weight gain was very significant (P<0.05) in Treatment-2 (P= 0.01), Treatment-4 (P=0.006), Control (P= 0.001) and Placebo (P=0.017) group but it was not significant in Treatment-1 (P= 0.09) and Treatment-3 (P= 0.07) group. On the other hand, during the repeated dose toxicity study, body weight also increased steadily but not significantly, as shown in **figure 1 (C)**. we saw those differences of body weight in all study groups in different evaluation time point (day 0, 3 and 7) were not significant (P>0.05). This result held true if the same data were analyzed independently per gender or evaluation time point. Despite these differences, it is possible to detect a significant increase of this parameter for all the groups during the study, translated into a normal evolution of body weight which constitutes an indicator of health for the animals and further substantiates the non-toxicity of the GBPD002 under both single and repeated dose toxicity study. The absence of negative effects on body weight gain is favorable for the evaluation of the substance under the study, since one of the primary clinical symptoms of stress or illness on this rat strain is precisely the decrease of body weight. Therefore, an increase in body weight is indirect evidence of non-toxicity for the substance under analysis.

**Figure 1.**
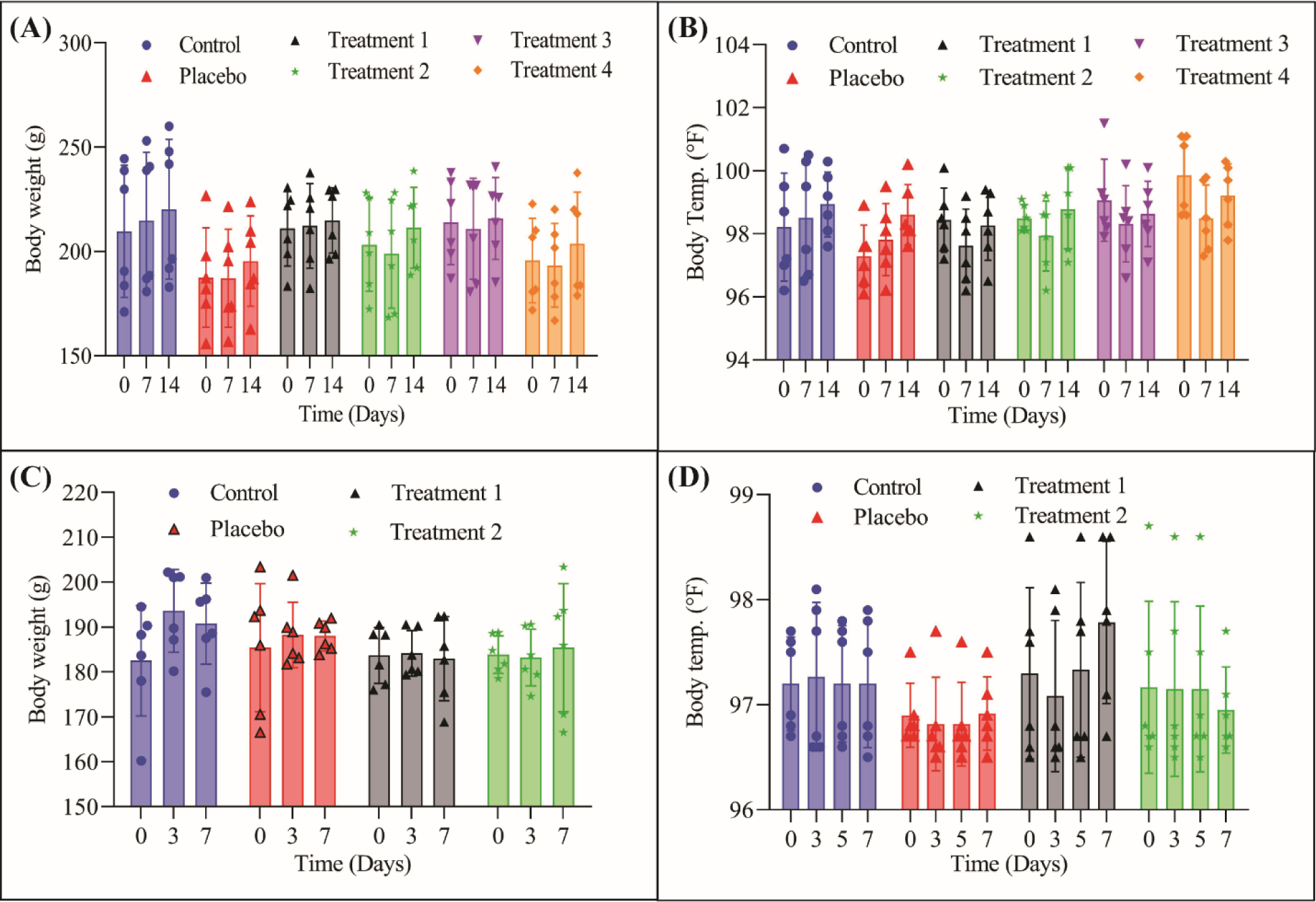
Behavior of body weight and temperature. (A) and (C) change of body weight for single and repeated dose toxicity study respectively. (B) and (D) change of body temperature for single and repeated dose toxicity study respectively.

### 3.2 Body temperature

During the single dose toxicity test, we found that the change of body temperature was not significant (P>0.05) in case of the 4 treatment groups (Treatment-1, Treatment-2, Treatment-3, Treatment-4) but it was more significant in Placebo (P= 0.009) and near significant in Control (P= 0.07), as shown in **figure 1 (B)**. On the other hand, it was observed that the change of body temperature was not significant (P>0.05) in case of all study groups (Treatment-1, Treatment-2, Placebo and Control), as shown in **figure 1 (D)**. This result also remained similar if the same data were analyzed independently per gender or evaluation time point which indicated normal health for the animals and further substantiates the non-toxicity of the GBPD002. However, no change in body temperature out of the normal range is an indirect indicator of non-toxicity for the test item under analysis.

### 3.3 Hematology parameters

When analyzing the results of the hematological tests, differences were detected among treatment groups of both doses of GBPD002 and Reference (Eprex^®^).

#### 3.3.1 Single dose toxicity study

The change of RBC count was significant (P<0.05) in Treatment-1 (P= 0.02), Treatment-3 (P=0.02), and Placebo (P=0.00) but not significant (P>0.05) in Treatment-2 (P= 0.16), Treatment-4 (P= 0.14) and Control (P= 0.43) as shown in **figure 2 (A)**. We observed a significant (P<0.05) change of WBC count in Treatment-1 (P= 0.05), Treatment-4 (P= 0.038), and Control (P= 0.001) group, however, insignificant (P>0.05) change found in Treatment-2 (P= 0.80), Treatment-3 (P= 0.80) and Placebo (P= 0.86) as shown in **figure 2 (B)**. The change of HGB count was significant (P<0.05) in Treatment-1 (P= 0.01), Treatment-3 (P=0.00), and Placebo (P=0.001) group but not significant (P>0.05) in Treatment-2 (P= 0.11), Treatment-4 (P= 0.05) and Control (P= 0.27) as shown in **figure 2 (C)**. The difference of PLT count between the groups by doing t test where we did not find any significant (P<0.05) differences among the groups; **figure 2 (D)**. The percentage of HCT was also observed where the change of HCT percentage was significant (P<0.05) in Treatment-1 (P= 0.01), Treatment-3 (P=0.02), and Placebo (P=0.00), but not significant (P>0.05) in Treatment-2 (P= 0.218), Control (P= 0.55) and near significant in Treatment-4 (P= 0.08) as shown in **figure 2 (E)**. we found that the change of MCV value was significant (P<0.05) in Treatment-1 (P= 0.02), Treatment-4 (P=0.01), and Placebo (P=0.02) but not significant (P>0.05) in Treatment-2 (P= 0.20), Control (P= 0.86) and near significant in Treatment-3 (P= 0.07) as shown in **figure 2 (F)**. The amount of MCH was also measured which was significant (P<0.05) in Placebo (P=0.03) but not significant (P>0.05) in Treatment-1 (P= 0.54) Treatment-2 (P= 0.91), Treatment-3 (P= 0.37), Treatment-4 (P=0.97) and Control (P=0.91) as shown in **figure 2 (G)**. we found that the change of MCHC count was significant (P<0.05) in Treatment-1 (P= 0.03) and near significant in Placebo (P=0.06)), but not significant (P>0.05) in Treatment-2 (P= 0.47), Treatment-3 (P=0.28), Treatment-4 (P=0.40) and Control (P= 0.90). Here, initial and final value both were very close as shown in **figure 2 (H)**.

**Figure 2.**
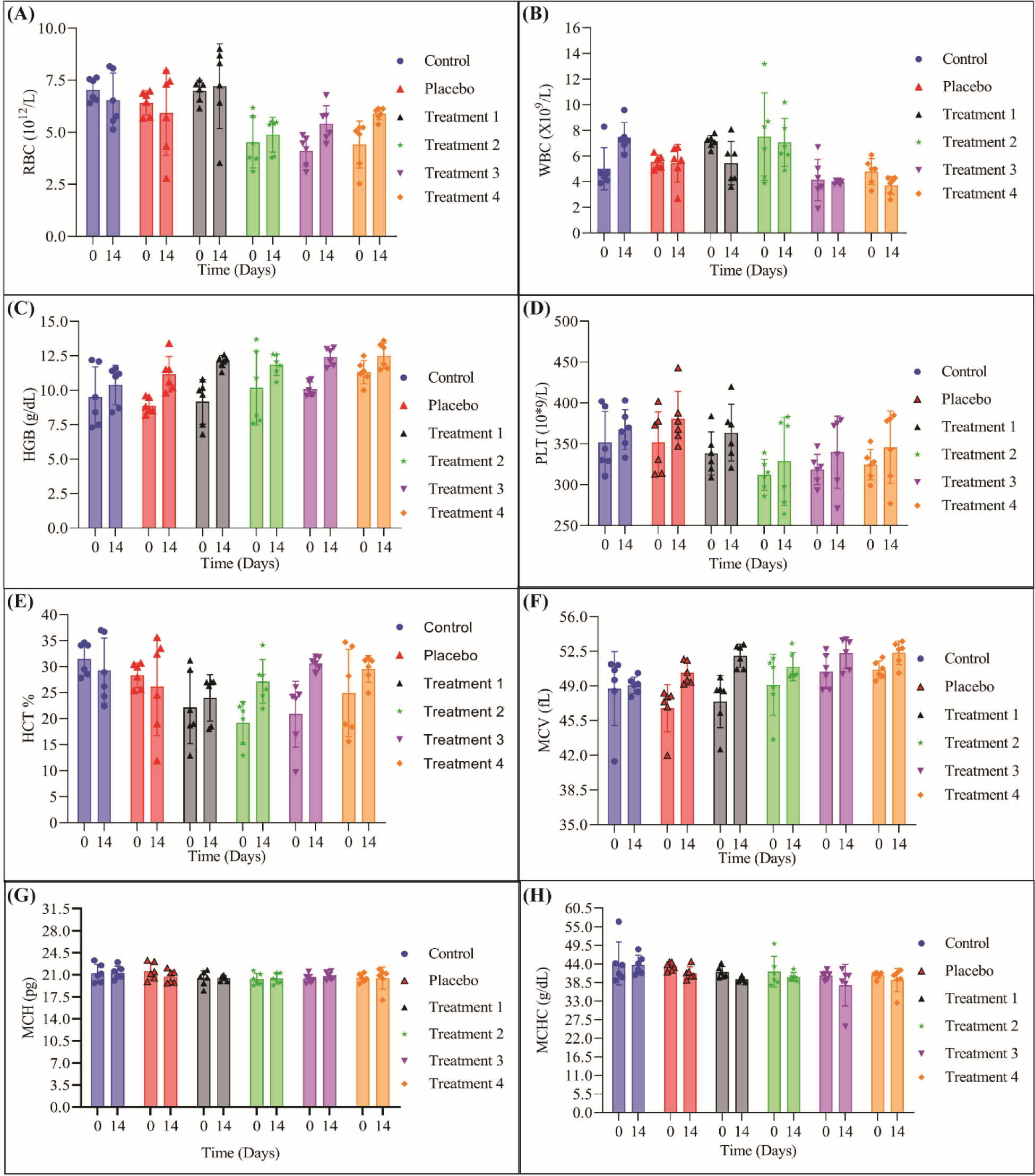
Hematology test parameters of single dose toxicity study. (A) red blood cell (RBC) count; (B) white blood cell (WBC) count; (C) hemoglobin count (HGB); (D) platelet (PLT) count; (E) hematocrit (HCT); (F) mean corpuscular volume (MCV); (G) mean corpuscular hemoglobin (MCH); (H) mean corpuscular hemoglobin concentration (MCHC).

#### 3.3.2 Repeated dose toxicity study

From one-way ANOVA and t test analysis the hematology parameters were observed on day-0 and Day-7 among the different study groups. RBC increased in all study groups but Treatment-1 and Treatment-2 showed highest count in comparison with Control and Placebo as shown in **figure 3(A)**. The differences of RBC count between the two treatment groups were very significant (P=0.0002). We also compared the differences between the groups by doing t test where we found significant differences between Treatment-1 and Control (P=0.002), Treatment-2 and Control (0.002), Treatment-1 and Placebo (P=0.002), Treatment-2 and Placebo (P=0.002) but not significant differences between Treatment-1 and Treatment-2 (P=0.83), and Placebo and Control (P=0.64). WBC decreased within normal range in all study groups including Control where the results were almost similar as shown in **figure 3(B)**. However, the differences of WBC count between the groups were analyzed where there was not any significance (P=0.80) in the differences. HGB increased in all study groups but Treatment-1 and Treatment-2 showed highest level in comparison with Control and Placebo as shown in **figure 3(C)**. However, differences of HGB level on day-0 and Day-7 between the different study groups were very significant (P=0.0004). HCT also increased in Treatment-1 and Treatment-2 groups but decreased in Control and Placebo as shown in **figure 3(E)**. The differences of HCT percentage between the different study groups were very significant (P=0.0001). It was also found the significant differences between Treatment-1 and Control (P=0.01), Treatment-2 and Control (0.000), Treatment-1 and Placebo (P=0.04), Treatment-2 and Placebo (P=0.000). MCV increased within normal range in all study groups including Control where Treatment-1 showed highest level in comparison with Treatment-2 Placebo and Control as shown in **figure 3(F)**. The differences of HCV count between the groups were observed where did not find any significance (P=0.64) in the differences. MCH decreased within normal range in all study groups except Treatment-2 where the results were almost similar (P=0.46) as shown in **figure 3(G)**. MCHC decreased in Treatment-2 and Control groups but increased in Treatment-1 and Placebo group as shown in **figure 3(H)**. The differences of MCH level between the groups were analyzed where it was not significant (P=0.27). Finally, PLT increased within normal range in all study groups including Control where the results were almost similar as shown in **figure 3(D)**. However, the differences of PLT count between the groups were found significant (P=0.44) in the differences.

**Figure 3.**
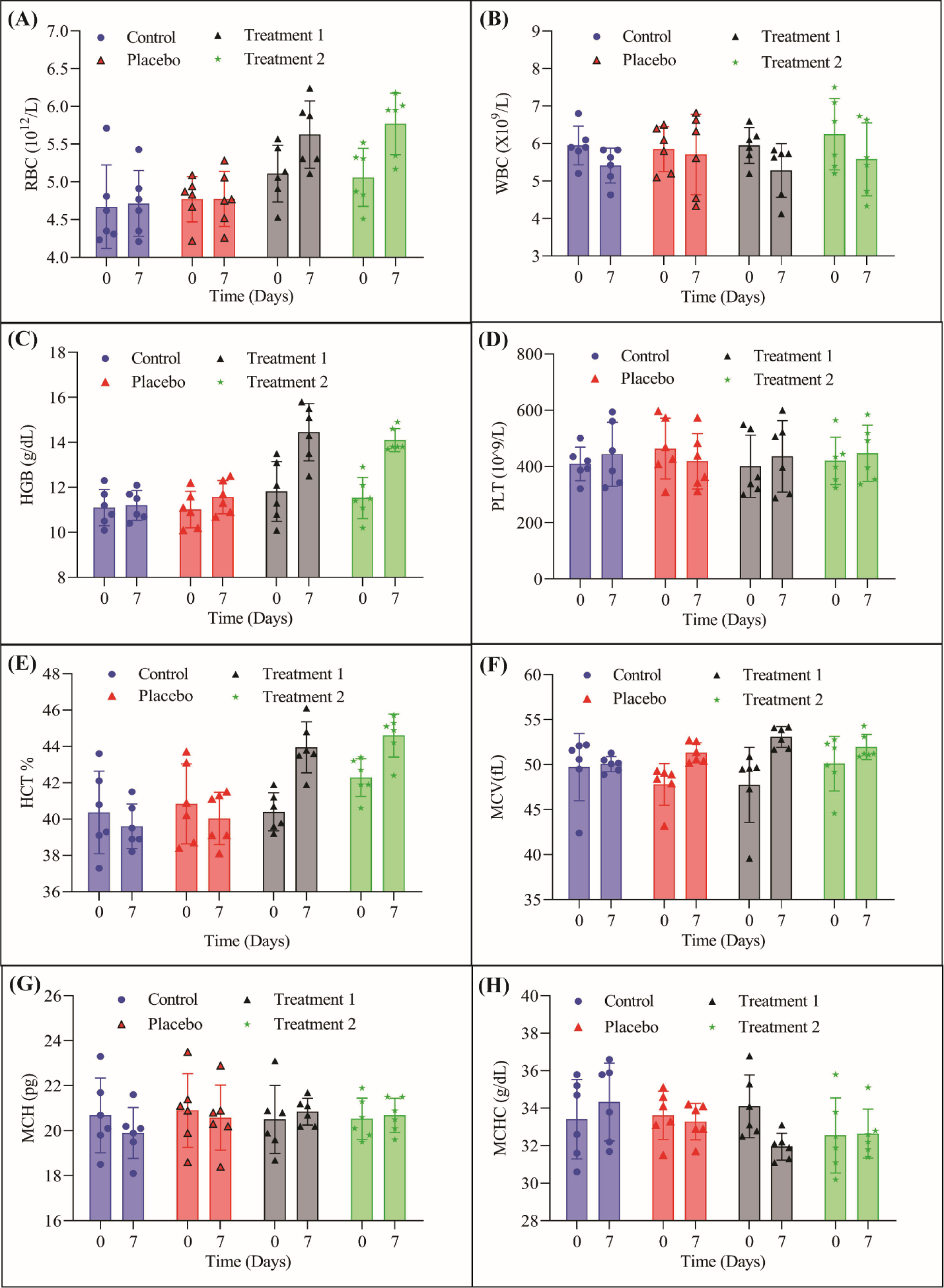
Hematology test parameters of repeated dose toxicity study. (A) red blood cell (RBC) count; (B) white blood cell (WBC) count; (C) hemoglobin count (HGB); (D) platelet (PLT) count; (E) hematocrit (HCT); (F) mean corpuscular volume (MCV); (G) mean corpuscular hemoglobin (MCH); (H) mean corpuscular hemoglobin concentration (MCHC).

### 3.4 Biochemical parameters test

#### 3.4.1 ALT/GPT

Increased ALT was observed in all animal group of the toxicity study. In single dose toxicity study, Treatment-4 group showed highest positive value among all the study groups, where Treatment-1 showed lowest result as shown in **figure 4 (A)**. The change of ALT was significant (P<0.05) in Treatment-2 (P= 0.03), Treatment-4 (P=0.00), Control (P= 0.01) and Placebo (P=0.06) group but not significant (P>0.05) in Treatment-1 (P= 0.14), and Treatment-3 (P= 0.10). On the other hand, in repeated dose toxicity test, ALT was also increased in all study groups where Treatment-2 and Control showed more increased result than Treatment-1 and Placebo as shown in **figure 4 (D)**. But the difference of ALT level between the treatment groups was not found any significant (P<0.05) differences among the groups.

**Figure 4.**
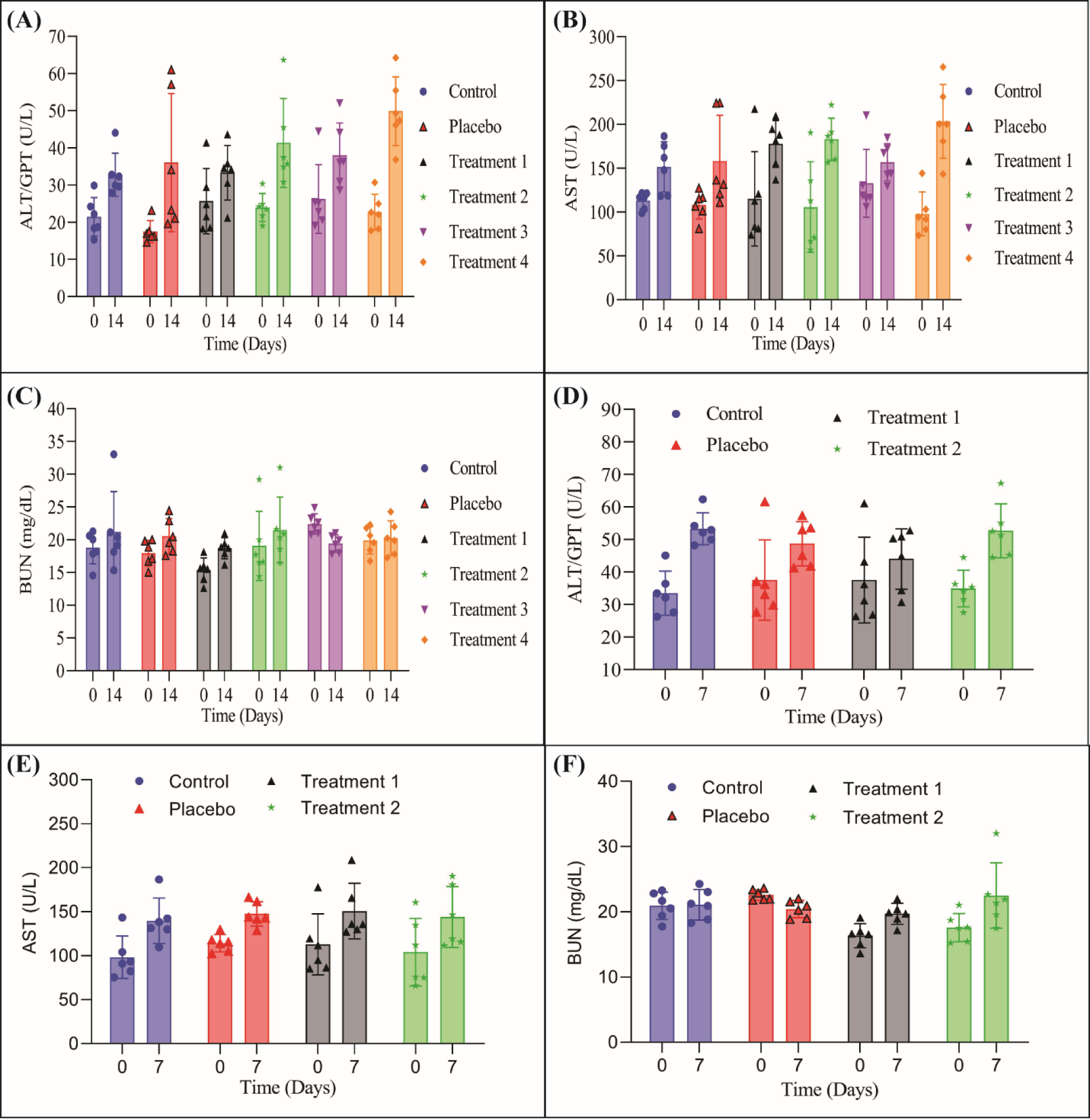
Analysis of biochemical parameters. (A), (B) and (C) for ALT/GPT, AST/GOT and BUN respectively for single dose toxicity test. (D), (E) and (F) for ALT/GPT, AST/GOT and BUN respectively for repeated dose toxicity test.

#### 3.4.2 AST

AST level increased positively in all animal groups under study where Treatment-4 showed more positive value than all other groups in the single dose toxicity study as shown in **figure 4 (B)**. The change of AST level was significant (P<0.05) in Treatment-1 (P= 0.02), Treatment-2 (P=0.02), Treatment-4 (P= 0.01), Control (P= 0.01) and Placebo (P=0.06) but not significant (P>0.05) in Treatment-3 (P= 0.28). The AST increased within normal range in all study groups including Control where the results were almost similar as shown in **figure 4 (E)**. However, the differences of AST level between the groups were analyzed where we did not find any significance (P=0.76) in the differences.

#### 3.4.3 BUN

BUN level also increased in all group under toxicity study except Treatment-3 in single dose toxicity test as shown in **figure 4 (C)**. The change of BUN level was significant (P<0.05) in Treatment-1 (P= 0.03), Treatment-2 (P=0.00), and Treatment-3 (P= 0.00) but not significant (P>0.05) in Treatment-4 (P= 0.80), and Control (P= 0.31) and Placebo (P=0.12). Here, initial and final value of both were very close. The BUN level also increased in Treatment-1, Treatment-2 and slightly in Control groups but decreased in Placebo in repeated dose toxicity study as shown in **figure 4 (F)**. However, the differences of BUN level between the groups were analyzed where it was not found any significance (P=0.02) in the differences.

### 3.5 Necropsy findings

Single representative subject from all treatment, placebo and control groups were euthanized for necropsy findings. External surfaces, all orifices, cranial cavities, external surfaces of brain and spinal cord, thoracic abdominal and pelvic cavities, cervical tissues and organs were observed for any abnormalities. However, any abnormal lesions were not found during necropsy examination as shown a **figure 5**.

**Figure 5.**
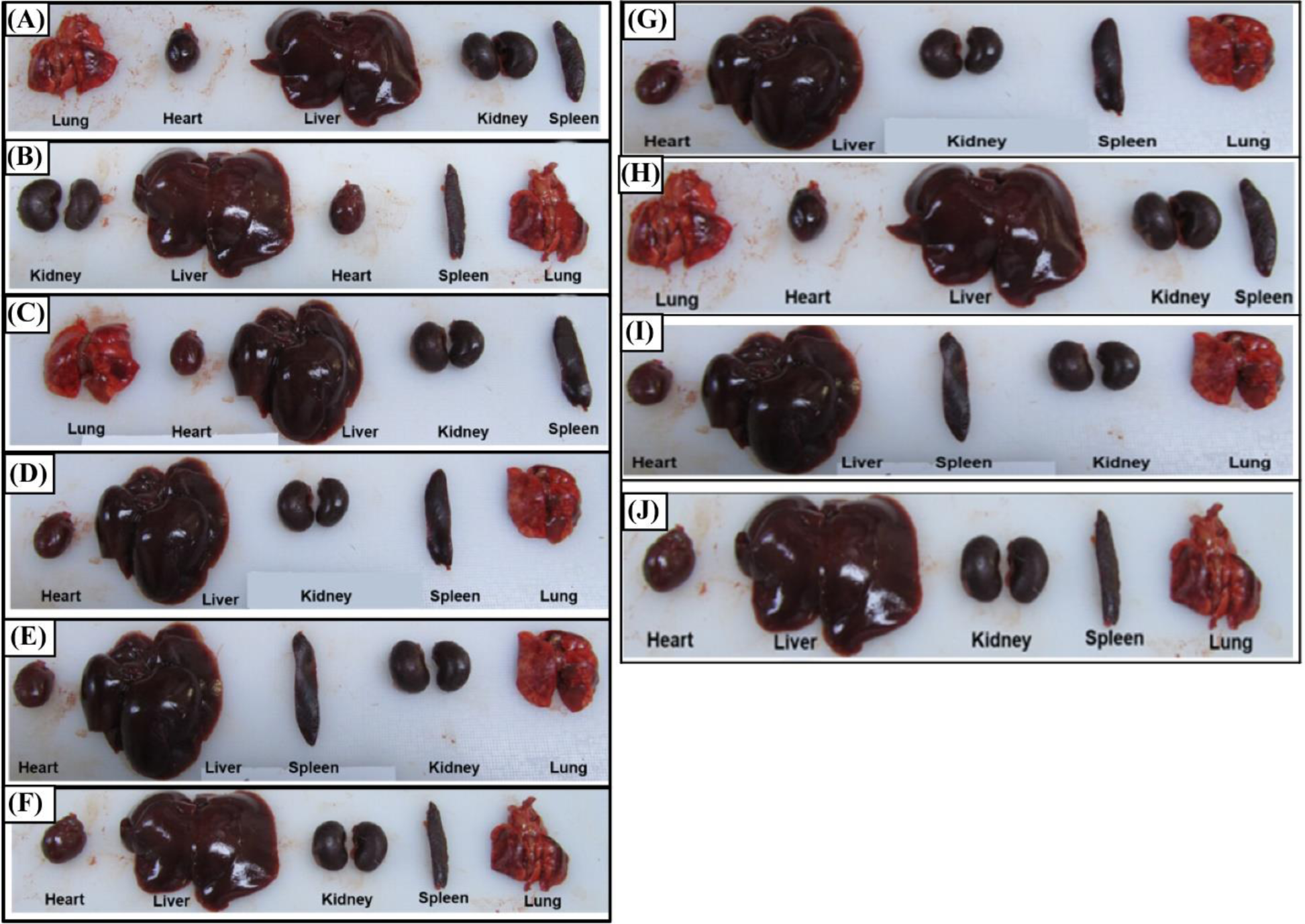
Necropsy findings of both single and repeated dose toxicity study. (A) Control, (B) Placebo, (C) Treatment 1, (D) Treatment 2, (E) Treatment 3, and (F) Treatment 4 groups of single dose toxicity study. (G) Control, (H) Placebo, (I) Treatment 1, and (J) Treatment 2 groups of repeated dose toxicity study.

### 3.6 Histopathology study

No morphological signs of toxicity were observed on internal organs such as kidney, liver, lung and spleen of any animals under both of single and repeated dose toxicity study (**Figure 6**). No lesion of pathological significance like degeneration or cellular necrosis were found on these internal organs among all treatment groups including the group which received Eprex^®^.

**Figure 6.**
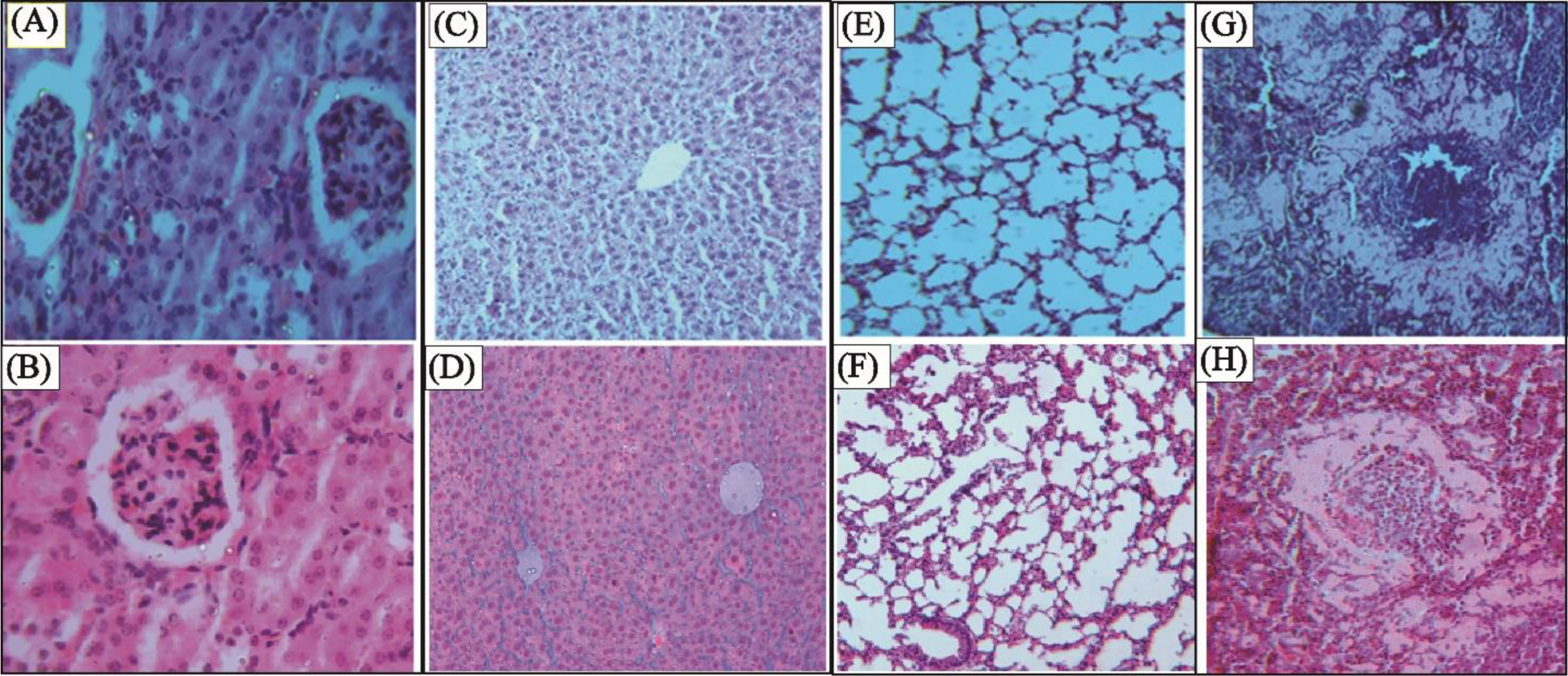
Histopathological findings. (A) and (B) kidney, (C) and (D) liver, (E) and (F) lung, (G) and (H) spleen both single and repeated dose toxicity study respectively.

## 4. Discussion

This study was reported the overall results of the preclinical evaluation of the safety of GBPD002 in the rat through a comparative study of acute toxicity with Eprex^®^, following subcutaneous administration. The study clearly established that the toxicological profile of GBPD002 and Eprex^®^, were similar and related to the known pharmacology of erythropoietin alfa previously used for a comprehensive safety and toxicity analysis of rhEPO [24]. No side effects or signs or toxicity were observed after daily observations of the animals inoculated with GBPD002 or Eprex^®^ in the comparative acute toxicity assay. There were no changes in fur or pigmentation and the eyes and mucosal surfaces appeared normal, as did the somatomotor activity and behavior. Proper responses to stimulation were obtained, and no deaths were reported during the study. Upon evaluation of the inoculation site, no signs of damage attributable to the administration of rhEPO were evidenced [25-27]. The only abnormalities detected were clinical signs (hemorrhagic areas) that appeared on the repeated dose toxicity study, but these were related to the method and the reiterative nature of the inoculation rather than to the use of rhEPO itself, since they also appeared with the same intensity in the placebo group [28-30]. The results of the hematological tests were normal when compared to their values before the start of the study.

Body weight increased steadily and significantly (*P*<0.05) during the single dose toxicity study but we saw those differences of body weight in all study groups in different evaluation time point (day 0, 3 and 7) were not significant (*P*>0.05) during the repeated dose toxicity study. It is possible to detect a significant increase of this parameter for all the groups during the study, translated into a normal evolution of body weight for both genders which constitutes an indicator of health for the animals and further substantiates the non-toxicity of the GBPD002 under both the study. The absence of negative effects on body weight gain is favorable for the evaluation of the substance under study, since one of the primary clinical symptoms of stress or illness on this rat strain is precisely the decrease of body weight [31, 32]. Therefore, an increase in body weight is indirect evidence of non-toxicity for the substance under analysis. There was no significant change in body temperature observed among treatment groups. This result also remained similar if the same data were analyzed independently per gender or evaluation time point which indicated normal health for the animals and further substantiates the non-toxicity of the GBPD002. Studies, they were shown no change in body temperature out of the normal range is an indirect indicator of non-toxicity for the test item under analysis [33, 34].

The results of the hematological tests were normal when compared to their values before the start of the study. The differences were found in the groups inoculated with GBPD002, Eprex^®^, or with the placebo in both single and repeated dose. The values stayed within the normal range for the species used in the assay [28, 35]. WBC decreased within normal range in all study groups including control where the results were almost similar. No changes in WBC count out of the normal range is evidence of non-toxicity for the test item under analysis [36]. The high counts of RBC, HB and the high HCT, together with the slight increase in PLT, confirm the therapeutic effect of EPO. These high counts of hematological parameters declare the therapeutical activity of GBPD002 compared with Eprex^®^. These results are related to the mobilization of hematopoietic progenitor cells to the peripheral blood, given the mechanism of action of the product under assay [37]. On the other hand, MCV increased in all groups under study where Treatment-1 showed more positive value for MCV than all other groups under toxicity study where MCV increased within normal range in all study groups of repeated dose toxicity including Control where Treatment-1 showed highest level in comparison with Treatment-2 and Placebo. The difference of MCV level between the groups where we did not find any significant (*P*<0.05) differences among the groups. MCH increased in Treatment-2, Treatment-3 and Control but decreased in Treatment-1, Treatment-4 and Placebo where MCH decreased within normal range (*P*=0.46) in all study groups except Treatment-2 where the results were almost similar in repeated dose toxicity study. We found that MCHC decreased in all animal group under single dose toxicity study where MCHC decreased in Treatment-2 and Control groups but increased in Treatment-1 and Placebo in repeated dose toxicity study. However, no change in MCV, MCH and MCHC level out of the normal range is evidence of non-toxicity for the test item under analysis [38]. Collectively, these results shown here demonstrated similar responses for experimental EPO preparations, viz., GBPD002 and Eprex^®^.

Our result is in accordance with the findings of other studies where a similar level of biochemical parameters (ALT/GPT, AST and BUN) [39]. In both of study, we reported that serum ATL/GPT increased in all the treatment groups. Along with serum AST and BUN also increased from the normal level but the increasing level of AST and BUN were less than the ALT/GPT level in both single and repeated dose toxicity test. Some of similar studies suggest that the normal high level of biochemical parameters indicates the uniformity of liver function [40]. It is also evidence of non-toxicity for the test item under analysis [41]. So, our observation suggests that the toxicity of GBPD002 was safe that was similar to the reference product Eprex^®^.

As all of the organs had normal macroscopic morphology, the results of the necropsies for both experiments showed no indication of any abnormalities or anatomical changes. Examining the inoculation site for two animals in the placebo group and three animals in the treatment group after two doses revealed hemorrhagic regions. Presumably, the trauma from several subcutaneous injections at the same location is connected to this symptom [6]. Thus, the macroscopic results support the clinical findings that there are no changes related to erythropoietin and that there are no signs of local irritation or injury. It is crucial to stress that there were no reports, regardless of the dosage or volume of administration, as the lack of macroscopic damage in the organs and tissues of the experimental animals is a crucial factor in determining the safety of the product being tested in both trials. One of the observed histopathological findings, no lesion of pathological significance like degeneration or cellular necrosis were found on internal organs in both studies. These reactions have been described in the literature as a consequence of the intense metabolic activity of this organ, and their spontaneous appearance has been reported for this animal species [42]. The fact that they are present in the control groups indicates that their appearance is not dependent on the effect of the different doses under study for the rhEPO in any of the assayed formulations. It is also important to remark that even in this aspect the histopathological results indicate non-toxicity effect of test product (GBPD002) and similar to reference product (Eprex^®^).

In conclusion, given the absence of toxicity in both assays and the similarity in the response profile to that reported for Eprex^®^ (a registered and internationally recognized commercial homologue), the results of the use of GBPD002 support the safety of our product. On the basis of the results of the experimental phase and the histopathological study, GBPD002 indicates that it is not a toxic agent and the biological response to the products GBPD002 and Eprex^®^ is essentially equivalent. Therefore, they can be considered biosimilar in pre-clinical settings, and likely be administered interchangeably.

## Funding

Globe Biotech Limited funded this study.

## Author contributions

Conceptualization and study plan: Kakon Nag, Naznin Sultana and Sitesh Chandra Bachar; Test product development, manufacturing and evaluation: Maksudur Rahman Khan, Samir Kumar, Md. Enamul Haq Sarker, Mohammad Mohiuddin, Uttam Barman, Rony Roy and Bipul Kumar Biswas; Data analysis and interpretation: Mohammad Mohiuddin; Quality management: Mohammad Mohiuddin and Rony Roy; Manuscript writing and editing: Kakon Nag, Naznin Sultana, Mohammad Mohiuddin and Sheik Rejaul Haq; All authors have read and agreed to the manuscript.

## Declaration of competing interests

The authors declare that they have no competing interests.

## Ethical statement

The study plan and procedures for experiments were approved by the internal ethical review board (IECB-PCS: Internal Ethical Clearance Board for Pre-Clinical Study) of Globe Biotech Limited, which is complied with the local and international regulation.

## Acknowledgements

The study was funded by Globe Biotech Limited. We thank Md. Harunur Rashid, the chairman of Globe Pharmaceuticals Group of Companies, Ahmed Hossain, Md. Mamunur Rashid, Md. Shahiduddin Alamgir and Abdullah Al Rashid, the directors of Globe Pharmaceuticals Group of Companies for their continuous support and encouragement. We also thank Dibakor Paul, Zahir Uddin Babor, GM Sajib Hassan, and Mijanur Rahman for their support for information and facility management system.

## Availability of data and materials

The data that support the findings of this study are available within the article and its Supplementary information file, or are available from the corresponding author upon reasonable request.

## Consent for publication

Not applicable.

## Notes

### Competing Interest Statement

The authors have declared no competing interest.

